# Tumor-derived hypoxic small extracellular vesicles promote endothelial cell migration and tube formation via ALS2/Rab5/β-catenin signaling

**DOI:** 10.1101/2024.02.02.578446

**Authors:** Patricio Silva, Nadia Hernández, Héctor Tapia, Belén Gaete-Ramírez, Tania Flores, Daniela Herrera, Albano Cáceres-Verschae, Manuel Varas-Godoy, Vicente A. Torres

## Abstract

Tumor hypoxia has been associated with cancer progression, angiogenesis, and metastasis via modifications in the release and cargo composition of extracellular vesicles secreted by tumor cells. Indeed, hypoxic extracellular vesicles are known to trigger a variety of angiogenic responses via different mechanisms. We recently showed that hypoxia promotes endosomal signaling in tumor cells via HIF-1α-dependent induction of the guanine exchange factor ALS2, which activates Rab5, leading to downstream events involved in cell migration and invasion. Since Rab5-dependent signaling is required for endothelial cell migration and angiogenesis, we explored the possibility that hypoxia promotes the release of small extracellular vesicles containing ALS2, which in turn activate Rab5 in recipient endothelial cells leading to pro-angiogenic properties. In doing so, we found that hypoxia promoted ALS2 expression and incorporation as cargo within small extracellular vesicles, leading to subsequent transfer to recipient endothelial cells, promoting cell migration, tube formation and downstream Rab5 activation. Consequently, ALS2-containing small extracellular vesicles increased early endosome size and number in recipient endothelial cells, which was followed by subsequent sequestration of components of the β-catenin destruction complex within endosomal compartments, leading to stabilization and nuclear localization of β-catenin. These events converged in the expression of β-catenin target genes involved in angiogenesis. Knockdown of ALS2 in donor-tumor cells, which precluded its incorporation into small extracellular vesicles, prevented Rab5-downstream events and endothelial cell responses, which depended on Rab5 activity and guanine exchange factor activity of ALS2. These findings indicate that vesicular ALS2, secreted in hypoxia, promotes endothelial cell events leading to angiogenesis. Finally, these events might explain how tumor angiogenesis proceeds in hypoxic conditions.

## INTRODUCTION

Hypoxia is a common condition of the tumor microenvironment, which develops as result of uncontrolled and rapid growth of solid tumors, along with aberrant angiogenesis (1). This condition leads to the adaptation of tumor cells, along with the re-programming of cellular metabolism, gene expression, protein homeostasis and release of extracellular factors (1-3). Hence, hypoxia has been associated with tumor aggressiveness, poor prognosis and metastasis, by mechanisms that remain to be defined (1, 4). Particularly, hypoxia is known to promote tumor angiogenesis by modulating the behavior of resident endothelial cells via both direct and indirect mechanisms (5). Besides the well-described direct effects of hypoxia in endothelial cells, indirect mediators of angiogenesis induced by hypoxia remain poorly understood. Increasing evidence indicates that extracellular vesicles (EVs) liberated by tumor cells induce endothelial cell events associated with angiogenesis (6-11) and, most importantly, that these events are further increased under hypoxic conditions (12-15). Specifically, EVs derived from hypoxic tumor cells, including multiple myeloma, oral squamous cell carcinoma, colorectal, leukemia, and lung cancer, among others, induce endothelial cell migration and angiogenesis *in vitro*, when compared with their counterpart released in normoxic conditions (15-19). The identity of cargo contained within hypoxic EVs and the consequences of such cargo in recipient endothelial cells remain to be elucidated.

EVs are membranous vesicles produced by different types of cells and derived from the plasma membrane (microvesicles: 50-1000) or endosomal compartments (exosomes: 50-150 nm). In this context, EVs such as exosomes and microvesicles, which are smaller than 200 nm and defined as small EVs (sEVs) by the Minimal Information for Studies of Extracellular Vesicles (MISEV), play an important role in cell communication, because they can transport a plethora of cargo between cells, including proteins, nucleic acids, lipids and small molecules that converge in the activation of signaling pathways in recipient cells (20, 21). In endothelial cells, tumor-derived EVs have been shown to activate different downstream signaling pathways associated with angiogenesis, including mTOR, ephrin reverse signaling, VEGF, and β-catenin (10, 16, 18, 22). Intriguingly, the effects that hypoxia and other conditions of the tumor microenvironment have on EVs composition and EVs-dependent activation of such signaling pathways, remain poorly understood.

We previously identified ALS2 as a novel hypoxia-inducible target, which is increased in different tumor cell types and that is associated with the activation of the endosomal protein Rab5, accounting for hypoxia-driven tumor cell migration and invasiveness (23, 24). Rab5 is a small GTPase that is not only involved in tumor cell signaling, but it is also required for a variety of endothelial cell responses, including cell adhesion, migration and angiogenesis *in vitro* (25, 26).

Intriguingly, the possibility EVs, liberated by tumor cells in response to hypoxia, harbor the capability of activating Rab5-dependent events in endothelial cells, has not been explored. In this study, we show that tumor cells release EVs under hypoxic conditions, which harbor the capacity of increasing angiogenic-related responses in endothelial cells, via vesicular transfer of ALS2, which in turns activates Rab5-dependent signaling in endothelial cells, by promoting β-catenin signaling. These findings open new avenues to understand the mechanisms whereby hypoxia and the tumor microenvironment modulate a critical response associated with tumor aggressiveness.

## RESULTS

### Small extracellular vesicles derived from hypoxic tumor cells induce cell migration, tube formation *in vitro* and β-catenin signaling in endothelial cells

To determine the effects of tumor-derived hypoxic small extracellular vesicles (sEVs) on endothelial cell function, sEVs were obtained from A549 lung carcinoma cell cultures, grown by 24 h under normoxic (21% O_2_) or hypoxic conditions (1% O_2_) in a sealed chamber. Conditioned media were obtained and sEVs fractions were prepared by differential centrifugation, as previously described (27). Particle analysis and further characterization revealed that hypoxia impacted neither sEV size distribution (Figure 1A, B), nor protein marker composition in comparison with sEVs obtained in normoxia (Figure 1C). Interestingly, hypoxic sEVs were found to promote endothelial cell migration and angiogenesis *in vitro*, since sEVs obtained in hypoxia, but not those obtained in normoxia, promoted EA.hy926 endothelial cell migration (Figure 1D) and endothelial tube formation in comparison with non-treated endothelial cells (Figure 1E), which is in agreement with previous studies showing that sEVs secreted by hypoxic tumor cells depict increased pro-angiogenic potential (17, 19).

**Figure 1.**
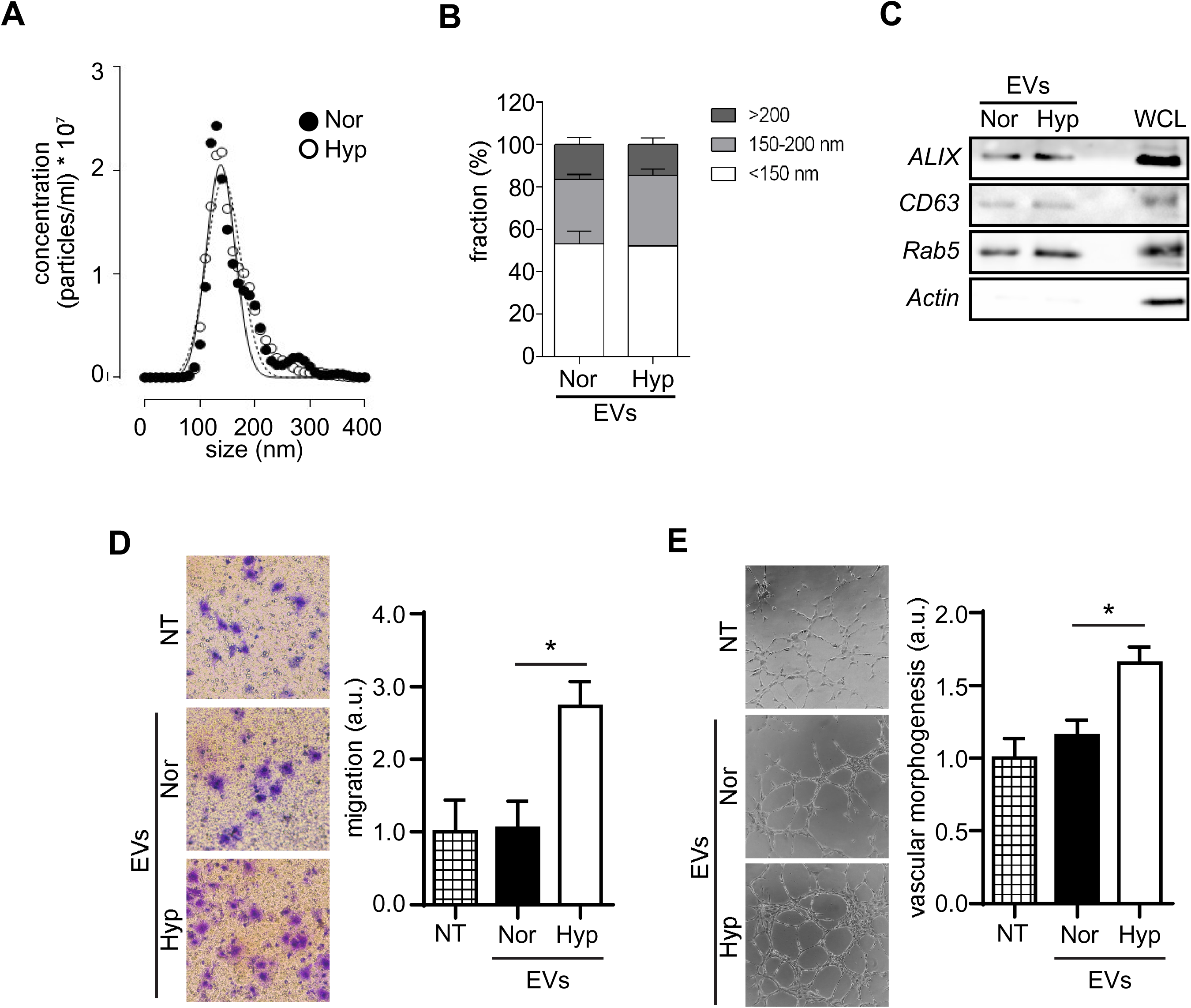
sEVs derived from hypoxic tumor cells induce endothelial cell migration and tube formation *in vitro*. **(A)** Particle size analysis. A549 cells were cultured in exosome-free medium, for 24h in normoxia (Nor) or hypoxia (Hyp). Conditioned media were obtained and sEV fractions were prepared by differential centrifugation. sEV fractions (100k pellets) were analyzed for size distribution and concentration in a NanoSight NS300 (Malvern). **(B)** Quantification of size distribution, obtained by averaging 3 independent purification experiments, as described in A (mean±s.e.m.). **(C)** Protein extracts were obtained from equal amounts of sEVs and samples were analyzed by Western blot for ALIX and CD63 (exosome markers), as well as Rab5 and Actin, using specific antibodies. Whole-cell lysates (WCL) obtained from A549 cells were included as a control. Images are representative from 3 independent experiments. **(D)** sEV fractions were added to normoxic EA.hy926 endothelial cells (1x10^4^ sEVs/cell), incubated for 24h, and then cells were harvested and allowed to migrate for 1.5h in Transwell chambers coated with 2 μg/ml fibronectin. Data were averaged from 4 independent experiments (mean±s.e.m.; *p<0.05). **(E)** sEVs fractions were added to normoxic EA.hy926 cells (1x10^4^ sEVs/cell), incubated for 24h and angiogenesis *in vitro* was measured by the endothelial tube formation assay. Data were averaged from 3 independent experiments (mean±s.e.m.; *p<0.05).

We next searched for signaling pathways that might be involved in sEV-driven effects in endothelial cells. Previous studies showed that sEVs promote endothelial cell responses via activation of Wnt/β-catenin signaling (18, 28), which is a pathway commonly involved in angiogenesis (29, 30), however, whether these effects are modulated by hypoxia, remain unknown. Hence, we evaluated whether hypoxia further enhances the effects of tumor cell-derived sEVs on β-catenin signaling in endothelial cells. First, immunofluorescence and subcellular fractionation assays showed that hypoxic sEVs substantially increased nuclear localization of β-catenin in endothelial cells, in comparison with normoxic sEVs (Figure 2A, B). In accordance with these observations, hypoxic sEVs further promoted β-catenin-Tcf/Lef-dependent transcription in recipient endothelial cells, as compared with cells left untreated or treated with normoxic sEVs (Figure 2C). The latter was associated with increased expression of the β-catenin-target genes MMP2 and VEGF-A, as determined by mRNA levels (Figure 2D). Next, we determined whether β-catenin signaling was required for sEVs-dependent effects in endothelial cells. First, by using the C59 compound, which was previously shown to prevent nuclear localization of β-catenin in epithelial cells ((31, 32), Supplementary Figure S1A), we determined whether hypoxic sEVs require β-catenin to promote endothelial cell migration, observing that C59 precluded hypoxic sEVs-driven endothelial cell migration (Figure 2E). These observations were further supported by targeting total levels of β-catenin, as siRNA-mediated knockdown of endogenous β-catenin prevented hypoxic sEVs-induced endothelial cell migration (Supplementary Figure S1B, Figure 2F). Taken together, these results indicate that β-catenin is relevant to the effects of hypoxic sEVs in endothelial cells.

**Figure 2.**
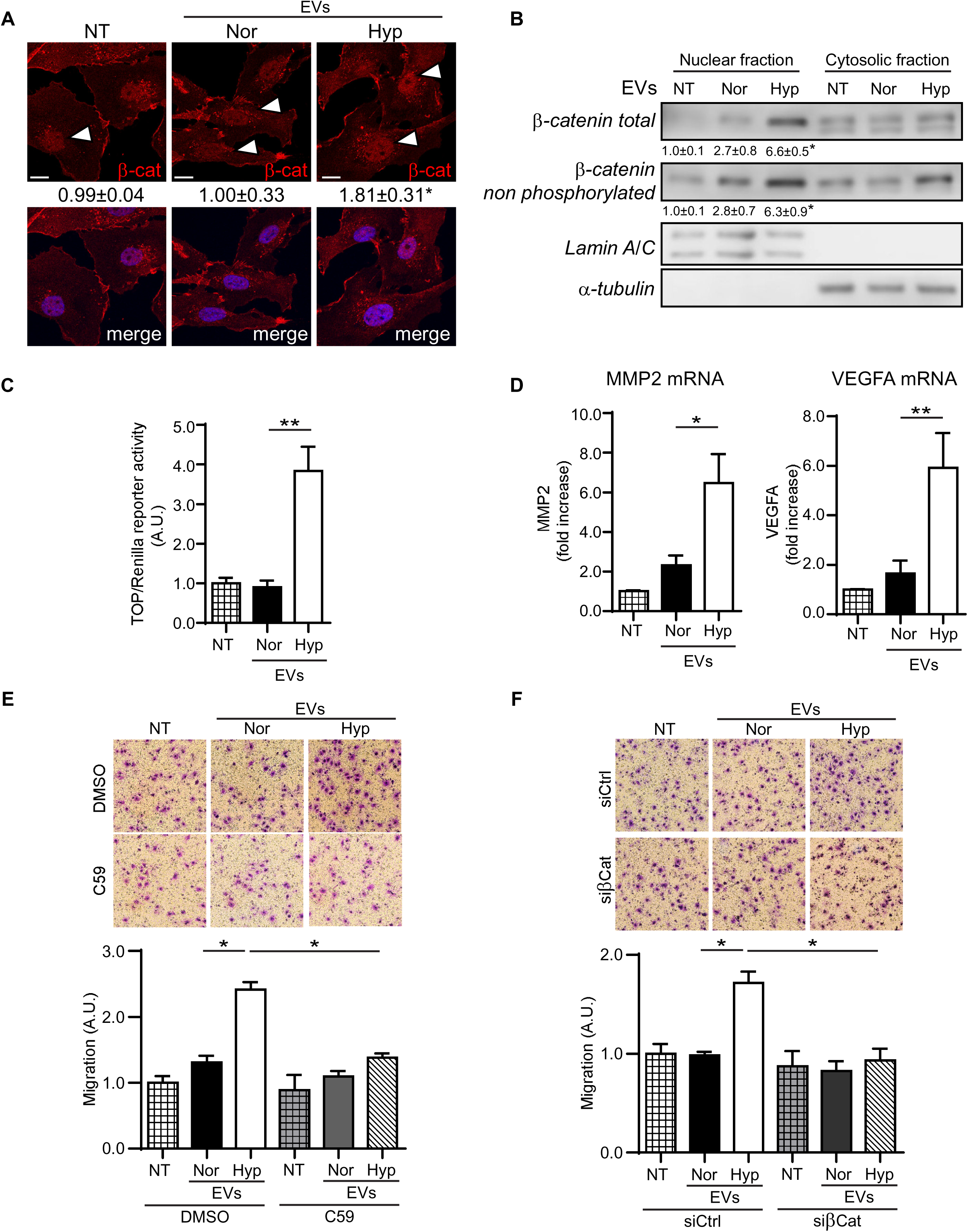
Hypoxic sEVs promote β-catenin signaling in endothelial cells. **(A)** sEV fractions were prepared by differential centrifugation (100k pellets) from conditioned media obtained from A549 cells that were incubated for 24h in normoxia (Nor) or hypoxia (Hyp). Then, sEVs were added to normoxic EA.hy926 endothelial cells (1x10^4^ sEVs/cell), incubated for 24h and samples were fixed and stained with an antibody against total β-catenin, whereas nuclei were visualized with Hoescht. Representative confocal microscope images are shown. Bar represents 10 µm. Numbers below each panel indicate normalized nuclear fluorescence signal, obtained by dividing nuclear versus total fluorescence signal, as described in the materials and methods section, using the Image J software. Data represent the average from 4 independent experiments (mean±s.e.m.; *p<0.05). **(B)** Subcellular fractionation of EA.hy926 cells left untreated (-) or treated with normoxic (Nor) or hypoxic (Hyp) sEVs for 24h. Total and non-phosphorylated (transcriptionally active) β-catenin were evaluated by Western Blot, using specific antibodies. Lamin A/C and α-tubulin were used as nuclear and cytosolic markers, respectively. Representative images are shown. Numeric data below each panel represent the scanning densitometric analysis obtained from 3 independent experiments (mean±s.e.m.; *p<0.05). **(C)** EA.hy926 were co-transfected with the plasmid pTOP-FLASH (functional Tcf/Lef binding sites, black bars) along with the constitutively active vector encoding for renilla luciferase (as control), and then treated or not with normoxic (Nor) or hypoxic (Hyp) sEVs. After 24 h, cells were homogenized and prepared for Tcf/Lef reporter assays (see materials and methods for details). Data were obtained as the ratio luciferase/renilla activity and shown as the relative values with respect to non-treated cells. Data were averaged from 4 independent experiments (mean±s.e.m.; **p<0.01). **(D)** EA.hy926 were stimulated for 24h with normoxic (Nor) or hypoxic (Hyp) sEVs, and then, total RNA was extracted, for subsequent RT-qPCR analysis of MMP2 and VEGFA. Relative abundance of all screened mRNAs was normalized to actin, by using the ΔΔCt method. Graphs show data obtained by averaging 4 independent experiments (mean±s.e.m.; *p<0.05; **p<0.01). **(E)** EA.hy296 cells were incubated or not with normoxic (Nor) or hypoxic (Hyp) sEVs for 24h, in the presence of vehicle (DMSO) or C59 (5 μM). Then, cells were harvested and allowed to migrate for 1.5h in Transwell chambers coated with 2 μg/ml fibronectin. Data represent the quantification of 3 independent experiments (mean ± s.e.m.; *p<0.05). **(F)** EA.hy296 cells were transfected with siRNA sequences targeting β-catenin or control and then incubated with normoxic (Nor) or hypoxic (Hyp) sEVs for 24h. Thereafeter, cells were harvested and allowed to migrate for 1.5h in Transwell chambers coated with 2 μg/ml fibronectin. Data represent the quantification of 4 independent experiments (mean ± s.e.m.; *p<0.05).

### Hypoxic sEVs promote endosomal sequestration of the β-catenin destruction complex

Former studies indicated that β-catenin stabilization, nuclear localization and transcription, proceed via a mechanism involving endosomal sequestration of the so-called “β-catenin destruction complex” (33). To explore this possibility, we assessed endosomal localization of components of the β-catenin destruction complex, including GSK3β, Axin and β-catenin itself, in a proteinase K/digitonin-resistance assay, as previously reported (32, 33). Remarkably, treatment of endothelial cells with hypoxic sEVs promoted resistance to digitonin/proteinase K in all components of the destruction complex (GSK3β, Axin, β-catenin), as compared with cells left untreated or treated with normoxic sEVs (Figure 3A, B). The latter was further confirmed by immunofluorescence analysis, which showed increased co-localization between Axin and the endosomal marker EEA1 (Figure 3C, D). The latter observations were intriguing, because previous studies indicated that endosomal localization of components of the β-catenin destruction complex is associated with fluctuations in early endosome dynamics (32). In fact, we found that hypoxic sEVs increase both the amount and size of EEA1-positive structures in recipient EA.hy926 cells (Figure 4A-C). Taken together, these data indicate that hypoxic sEVs promote endosomal sequestration of the β-catenin destruction complex, which is associated with fluctuations in EEA1-positive endosomes.

**Figure 3.**
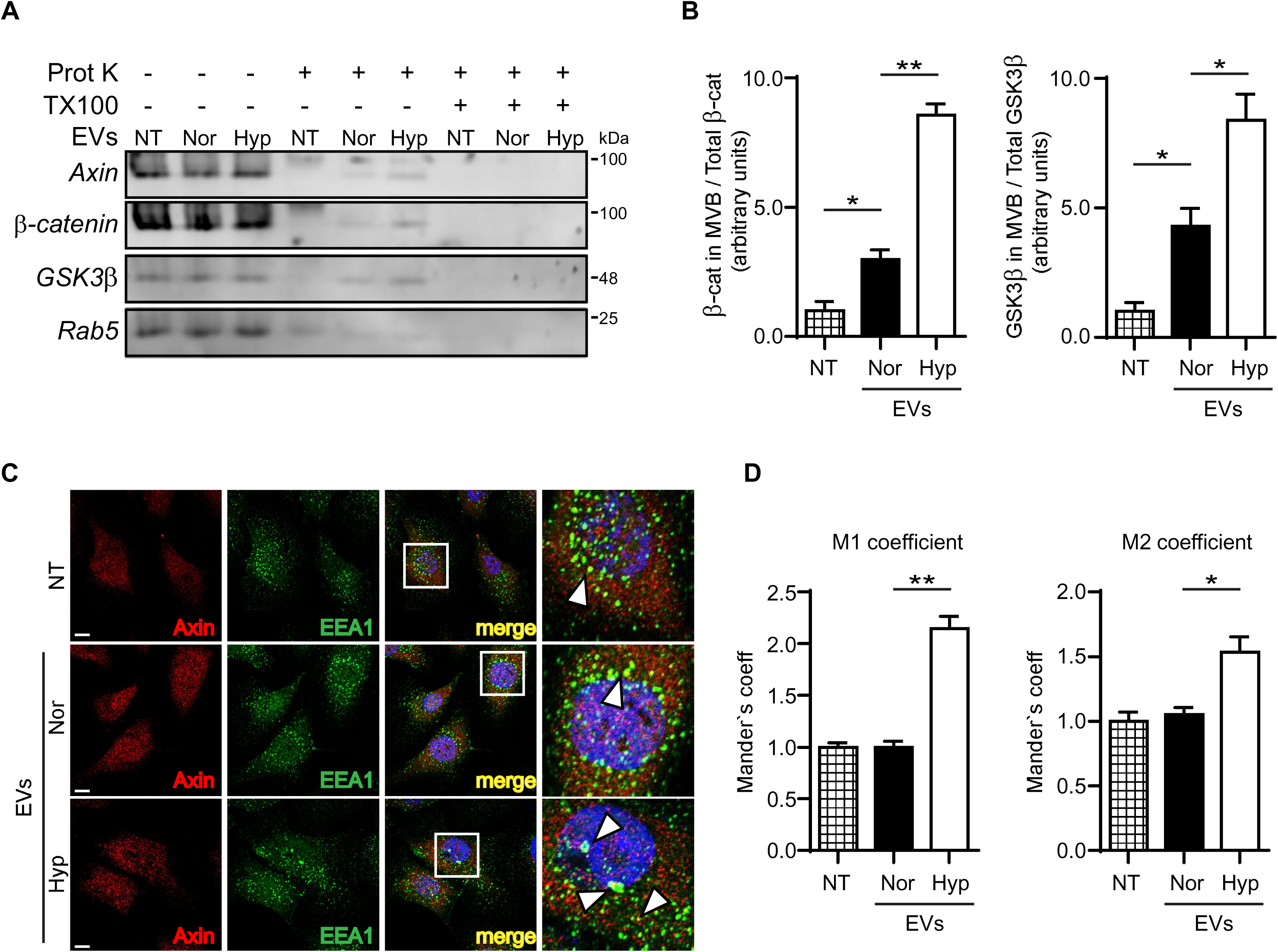
Hypoxic sEVs promote endosomal sequestration of the β-catenin destruction complex in endothelial cells. **(A)** sEV fractions were prepared by differential centrifugation (100k pellets) from conditioned media obtained from A549 cells that were incubated for 24h in normoxia (Nor) or hypoxia (Hyp). Then, sEVs were added to normoxic EA.hy926 endothelial cells (1x10^4^ sEVs/cell), incubated for 24h, and cells were harvested and processed for Proteinase K treatment, as described in the materials and methods. Samples were either left untreated or incubated with Proteinase K or Proteinase K + 0.1% Triton X-100, for 10 minutes. Samples were analyzed by Western blot, using antibodies raised against Axin, β-catenin, GSK3β or Rab5. Representative images are shown. **(B)** Graphs represent scanning densitometric data, obtained from (A) and averaged from 3 independent experiments (mean±s.e.m.; *p<0.05 **p<0.01). **(C)** EA.hy296 cells were left untreated or treated with normoxic (Nor) or hypoxic (Hyp) sEVs for 24 h, then processed for immunofluorescence analysis of EEA1 and Axin (EEA1, green; Axin, red; nuclei were stained with Hoescht, blue). Representative confocal microscopy images and their magnifications are shown from 3 independent experiments (Bar 10µm). **(D)** Mander’s coefficients were determined from data in (C), using JACOP plugins of Image-J software, as described in the material and methods section. Data were averaged from 3 independent experiments (mean±s.e.m; *p<0.05; **p<0.01).

**Figure 4.**
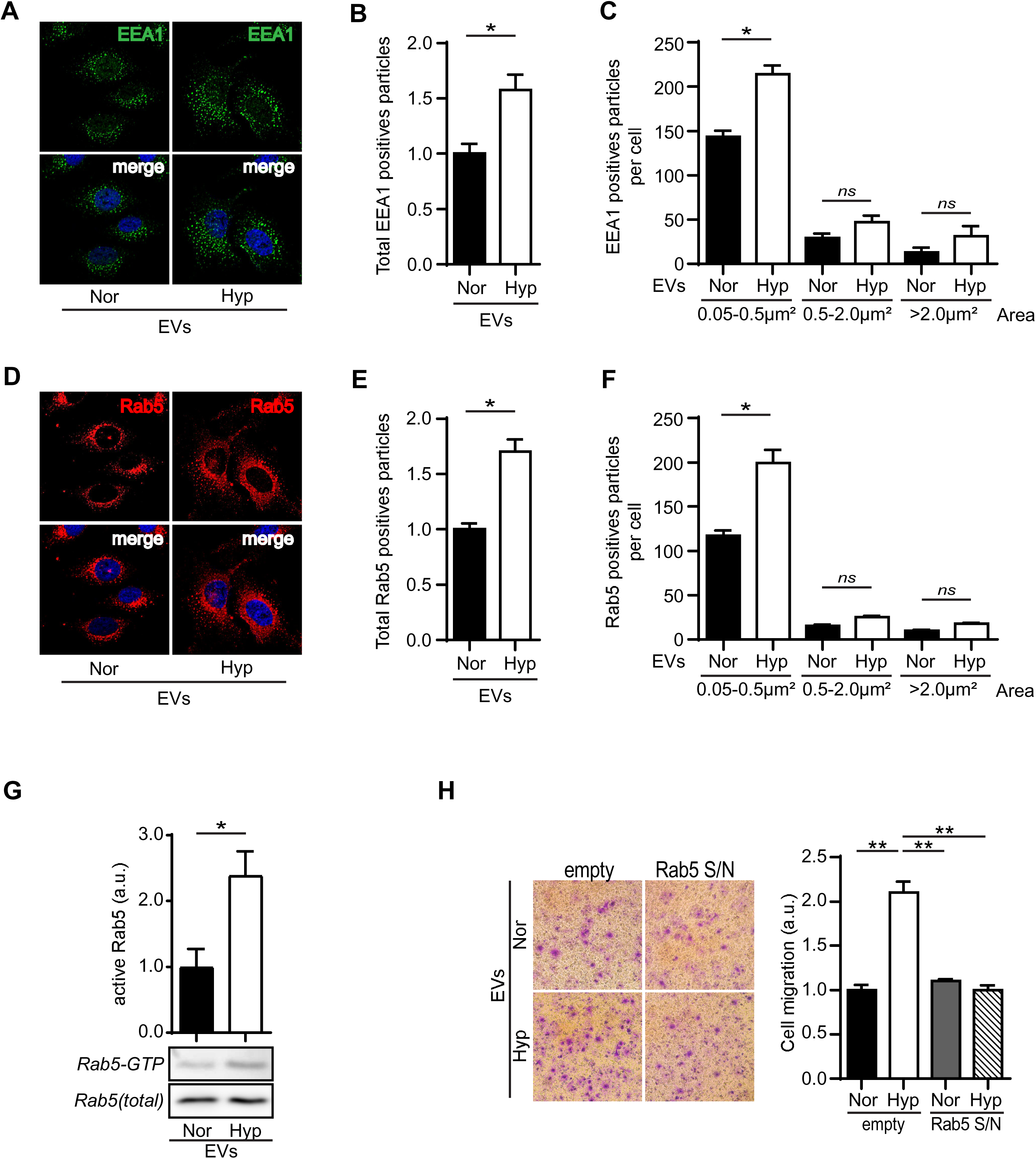
sEVs derived from hypoxic tumor cells promote Rab5 activation in endothelial cells. sEV fractions were prepared by differential centrifugation (100k pellets) from conditioned media obtained from A549 cells that were incubated for 24h in normoxia (Nor) or hypoxia (Hyp). Then, sEVs were added to normoxic EA.hy926 endothelial cells (1x10^4^ sEVs/cell), incubated for 24h, and samples were processed for subsequent analysis. **(A)** Immunofluorescence analysis of EEA1 (nuclei were stained with Hoechst). Representative confocal microscopy images are shown from 3 independent experiments (Bar 10µm). **(B)** Images described in (A) were analyzed for the total number of EEA1-positive particles per cell. Data represent the average of 3 independent experiments (mean±s.e.m.; **p<0.01). **(C)** EEA1-positive particles were grouped according to their size in three categories, 0.05-0.5, 0.5-2.0 and >2.0 μm2. At least 5 images per condition were analyzed using the Image J software, and the data represent the average of 3 independent experiments (mean±s.e.m.; *p<0.01). **(D-F)** Immunofluorescence analysis of Rab5 (nuclei were stained with Hoechst). **(D)** Representative confocal microscopy images are shown from 3 independent experiments (Bar 10µm). **(E)** Images described in (D) were analyzed for the total number of Rab5-positive particles per cell. Data represent the average of 3 independent experiments (mean±s.e.m.; **p<0.01). **(F)** Rab5-positive particles were grouped according to their size, as described in (C). Data represent the average of 3 independent experiments (mean±s.e.m.; *p<0.01). **(G)** EA.hy926 were stimulated for 24h with normoxic (Nor) or hypoxic (Hyp) sEVs, and then whole cell lysates were prepared for measurement of Rab5-GTP levels by pulldown. Images are representative and the graph shows the average of 5 independent experiments (mean±s.e.m.; *p<0.05). **(H)** EA.hy296 cells were transfected with the plasmids pEGFP-C1 (empty) or pEGFP-Rab5/S34N (Rab5 inactive mutant, GTPase deficient). Then, cells were incubated with sEVs (obtained as indicated in A) for 24 h, harvested, and therefore allowed to migrate for 1.5h in Transwell chambers coated with 2 μg/ml fibronectin. Data represent the quantification of 3 independent experiments (mean ± s.e.m.;**p<0.01).

Previous studies indicated that early endosome size and number were dependent on the small GTPase Rab5 (34), which is also known as a critical regulator of endothelial cell migration and angiogenesis *in vitro* (25, 26). Indeed, we observed that hypoxic sEVs increase both the amount and size of Rab5-positive structures in recipient EA.hy926 cells (Figure 4D-F), which was in agreement with data obtained for EEA1-positive structures (Figure 4A). Since Rab5 activity is required for endothelial cell responses and based on previous evidence indicating that Rab5 gets activated under hypoxic conditions (23, 24), we hypothesized that sEVs released by hypoxic tumor cells activate Rab5 in recipient endothelial cells, resulting in increased angiogenic-related responses. We first determined the effects normoxic and hypoxic sEVs on EAhy.926 cells, observing that hypoxic sEVs promote Rab5-GTP loading in recipient endothelial cells, as compared with those cells treated with normoxic sEVs (Figure 4G). Then, we determined the dependency of Rab5 activity in sEVs-induced endothelial cell migration, observing that hypoxic sEVs promoted endothelial cell migration in control (GFP-transfected) cells, but not in GFP-Rab5/S34N (the inactive mutant form of Rab5) transfected cells, indicating that Rab5 activity was required for the effects of hypoxic sEVs in endothelial cell migration (Figure 4H).

### Vesicular transfer of ALS2 is required for sEVs to induce Rab5 activation and endothelial cell responses

The observation that Rab5 gets activated in recipient endothelial cells treated with hypoxic sEVs and the dependency of Rab5 activity on sEVs-induced endothelial cell migration, prompted us to investigate the mechanisms underlying such activation. In this respect, previous studies by our group identified the Rab guanine exchange factor (GEF) ALS2 as responsible for Rab5 activation in hypoxic tumor cells (23). Remarkably, we found that tumor cell-derived sEVs contained ALS2 as cargo, and that this was further increased in hypoxia (Figure 5A, B). Thus, we explored the possibility that the effects of hypoxic sEVs on endothelial cells were dependent on vesicular ALS2. To this end, recipient EA.hy296 cells were transfected with plasmids encoding for either GFP alone (empty) or GFP-ALS2ΔVPS (a dominant negative form of ALS2, which lacks GEF activity towards Rab5, (35), followed by subsequent evaluation of sEVs-dependent endothelial cell migration. As anticipated, GFP alone did not affect the capacity of hypoxic sEVs to induce endothelial cell migration, however, these effects were abrogated in GFP-ALS2ΔVPS-transfected cells (Figure 5C). These data suggest that vesicular transfer of ALS2 accounts for sEVs-induced effects in endothelial cells and that these effects are associated with regulation of Rab5 activity. To further confirm the relevance of vesicular ALS2 on sEVs-dependent effects in recipient endothelial cells, we downregulated endogenous ALS2 in donor A549 tumor cells by shRNA and obtained sEVs in normoxic and hypoxic conditions. In accordance with previous studies, we found that ALS2 was upregulated by hypoxia in donor A549 cells, while shRNA-mediated targeting of endogenous ALS2 prevented hypoxia-dependent induction of ALS2 levels in tumor cells (Figure 5D). Next, we determined the levels of ALS2 in sEVs secreted by hypoxic tumor cells, finding that hypoxia increases vesicular ALS2 protein levels in shRNA-control treated cells, but not in shRNA-ALS2 treated cells (Figure 5E). Importantly, knockdown of ALS2 in donor cells did not affect sEVs size distribution or protein markers (Supplementary Figure S2, Figure 5E). Then, normoxic and hypoxic sEVs obtained from control or ALS2-downregulated cells were used for subsequent analyses of sEVs-induced effects in endothelial cells, finding that vesicular ALS2 was required for Rab5-GTP loading (Figure 5F) and increases of early endosome size and number (Figure 5G).

**Figure 5.**
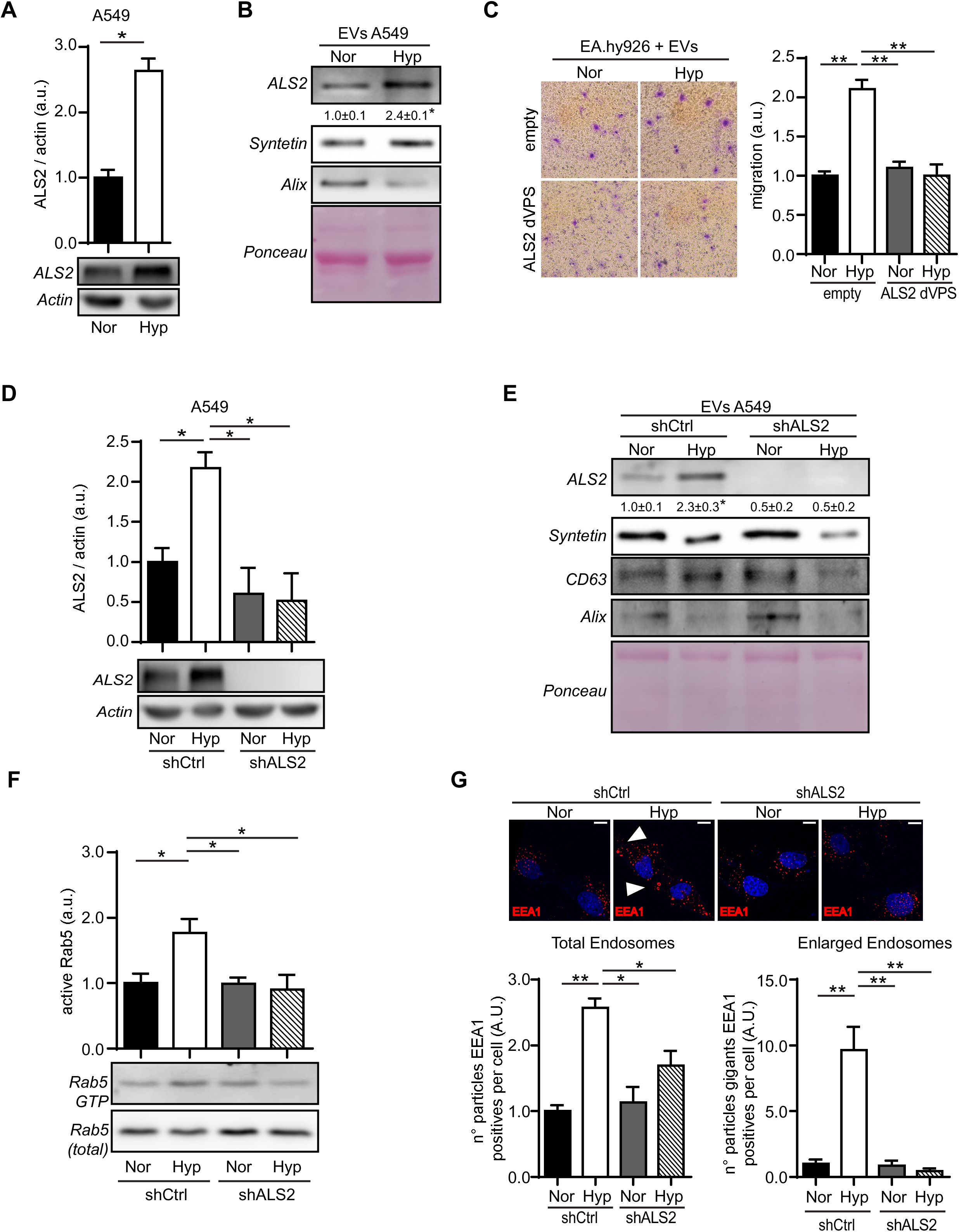
Extracellular vesicular ALS2 is associated with sEV-induced cell migration, Rab5 activation and endosome dynamics in endothelial cells. **(A)** A549 cells were grown in normoxia (Nor) or hypoxia (Hyp) for 24h and whole cell lysates were prepared for subsequent Western blot analysis of ALS2 and Actin. Images are representative and the graph shows the average of 3 independent experiments (mean±s.e.m.; *p<0.05). **(B)** sEV fractions were prepared by differential centrifugation (100k pellets) from conditioned media obtained from A549 cells cultured in normoxia or hypoxia, as described in (A), and protein extracts were prepared from equal amounts of sEVs. Samples were analyzed by Western blot of ALS2 and the exosomal markers Syntenin and Alix, as well as for Ponceau red staining. Images are representatives of 3 independent experiments. Numerical data below panels indicate the quantification obtained by scanning densitometry. (mean±s.e.m.;*p<0.05). **(C)** EA.hy296 cells were transfected with the plasmids pEGFP-C1 (empty) or pEGFP-ALS2ΔVPS (ALS2 dominant negative, lacking Rab5-GEF activity). Then, cells were incubated with sEVs (obtained as indicated in B) for 24 h, harvested, and therefore allowed to migrate for 1.5h in Transwell chambers coated with 2 μg/ml fibronectin. Data represent the quantification of 3 independent experiments (mean ± s.e.m.;**P<0.01). **(D)** A549 cells were treated with shRNA sequences against human ALS2 (scramble sequence was used as control) and selected for stable expression. Then, cells were grown for 24h in normoxia (Nor) or hypoxia (Hyp) and whole cell lysates were prepared for subsequent Western blot analysis of ALS2 and Actin. Images are representative and the graph shows the average of 3 independent experiments (mean±s.e.m.; *p<0.05). **(E)** sEV fractions were prepared as described in (B), from shRNA-control and shRNA-ALS2 cells obtained in (D). Protein extracts were prepared from equal amounts of sEVs and analyzed by Western blot of ALS2 and the exosomal markers Syntenin, Alix and CD63. Ponceau red staining is also shown. Images are representatives of 3 independent experiments. Numerical data below panels indicate the quantification obtained by scanning densitometry. (mean±s.e.m.; *p<0.05). **(F)** sEV fractions prepared in (E) were added to normoxic EA.hy926 cells (1x10^4^ sEVs/cell), incubated for 24h and Rab5-GTP levels were measured by pulldown assay. Images and the graph show the average of 5 independent experiments (mean±s.e.m.; *p<0.05). **(G)** EA.hy926 cells were treated with sEVs (obtained as indicated in B) for 24 h and processed for subsequent immunofluorescence analysis of EEA1 (nuclei were stained with Hoechst). Representative confocal microscopy images are shown from 3 independent experiments (Bar 10µm). The graphs show the amount of total and enlarged EEA1-positive particles per cell, as the average of 3 independent experiments (mean±s.e.m.; *p<0.05; **p<0.01).

By looking at the role of vesicular ALS2 in endothelial cell events, we found that shRNA-mediated downregulation of ALS2 in tumor cells precluded hypoxic sEVs-induced endothelial cell migration (Figure 6A) and tube formation *in vitro* (Figure 6B), which supported our previous findings (Figure 1D, E and Figure 5F). Collectively, these data indicate that extracellular vesicular ALS2, induced in hypoxia, promotes the activation of Rab5 in recipient endothelial cells and that this activation is important for the functional effects triggered by hypoxic sEVs in endothelial cells.

**Figure 6.**
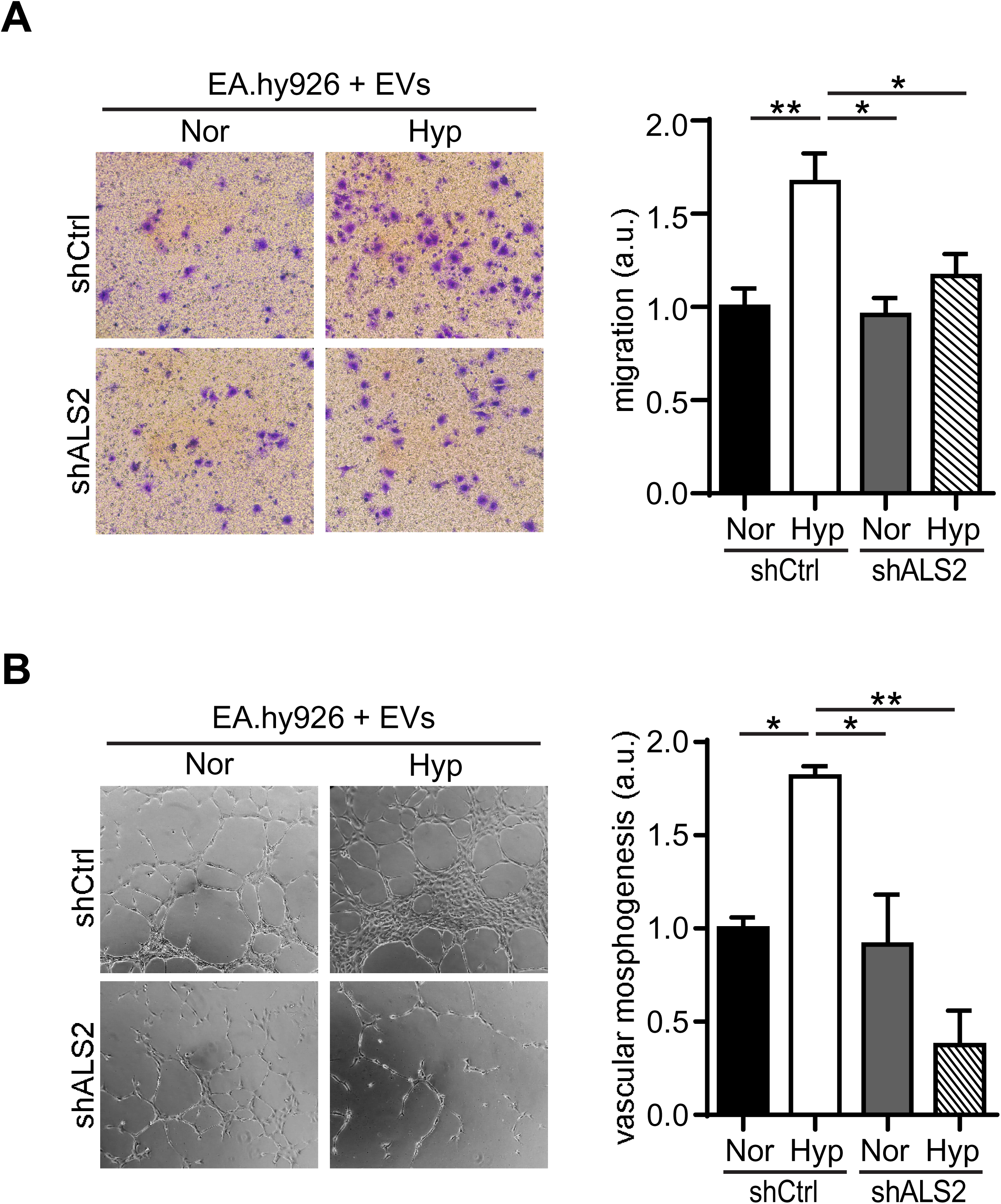
Extracellular vesicular ALS2 is required for hypoxic sEVs-induced endothelial cell migration and tube formation. sEV fractions were prepared by differential centrifugation (100k pellets) from conditioned media obtained from A549 cells that were incubated for 24h in normoxia (Nor) or hypoxia (Hyp). Then, sEVs were added to normoxic EA.hy926 endothelial cells (1x10^4^ sEVs/cell), incubated for 24h, and samples were processed for subsequent analysis. **(A)** Transwell migration assays were performed for 1,5h, using chambers coated with 2 μg/ml fibronectin. Data represent the quantification of 4 independent experiments (mean±s.e.m.; *p<0.05; **p<0.01). **(I)** Angiogenesis *in vitro* was measured by the endothelial tube formation assay. Data were averaged from 3 independent experiments (mean±s.e.m.; *p<0.05; **p<0.01).

### Role of ALS2 in sEVs-induced β-catenin signaling in endothelial cells

Next, we explored the functional consequences of downregulating extracellular vesicular ALS2 in β-catenin signaling, as a pro-angiogenic pathway activated downstream of Rab5 (32). First and in agreement with our early results (Figure 2), sEVs obtained from hypoxic tumor cells treated with a shRNA-control promoted nuclear localization of β-catenin in recipient endothelial cells, however, these events were abolished following downregulation of ALS2 in sEVs (Figure 7A). Moreover, sEVs obtained from hypoxic shRNA-control cells, but not shRNA-ALS2 targeted cells, promoted β-catenin-Tcf/Lef-dependent transcription, further indicating that vesicular ALS2 is required for β-catenin signaling in recipient endothelial cells (Figure 7B). These observations were further supported by vesicular ALS2-dependent expression of the β-catenin target genes MMP2 and VEGFA (Figure 7C). Interestingly, we found that hypoxic sEVs also promoted endosomal localization of β-catenin in recipient endothelial cells, which is in agreement with the proteinase K protection data (Figure 3), however, downregulation of ALS2 within hypoxic sEVs prevented this endosomal localization (Figure 7D, E). Interestingly and in agreement with our previous data, we noted that recipient endothelial cells treated with control hypoxic sEVs, but not ALS2-downregulated sEVs, showed the enlargement of EEA1-positive early endosomes (Figure 7D). Taken together, these data indicate that ALS2 contained within hypoxic sEVs promotes β-catenin signaling in recipient endothelial cells.

**Figure 7.**
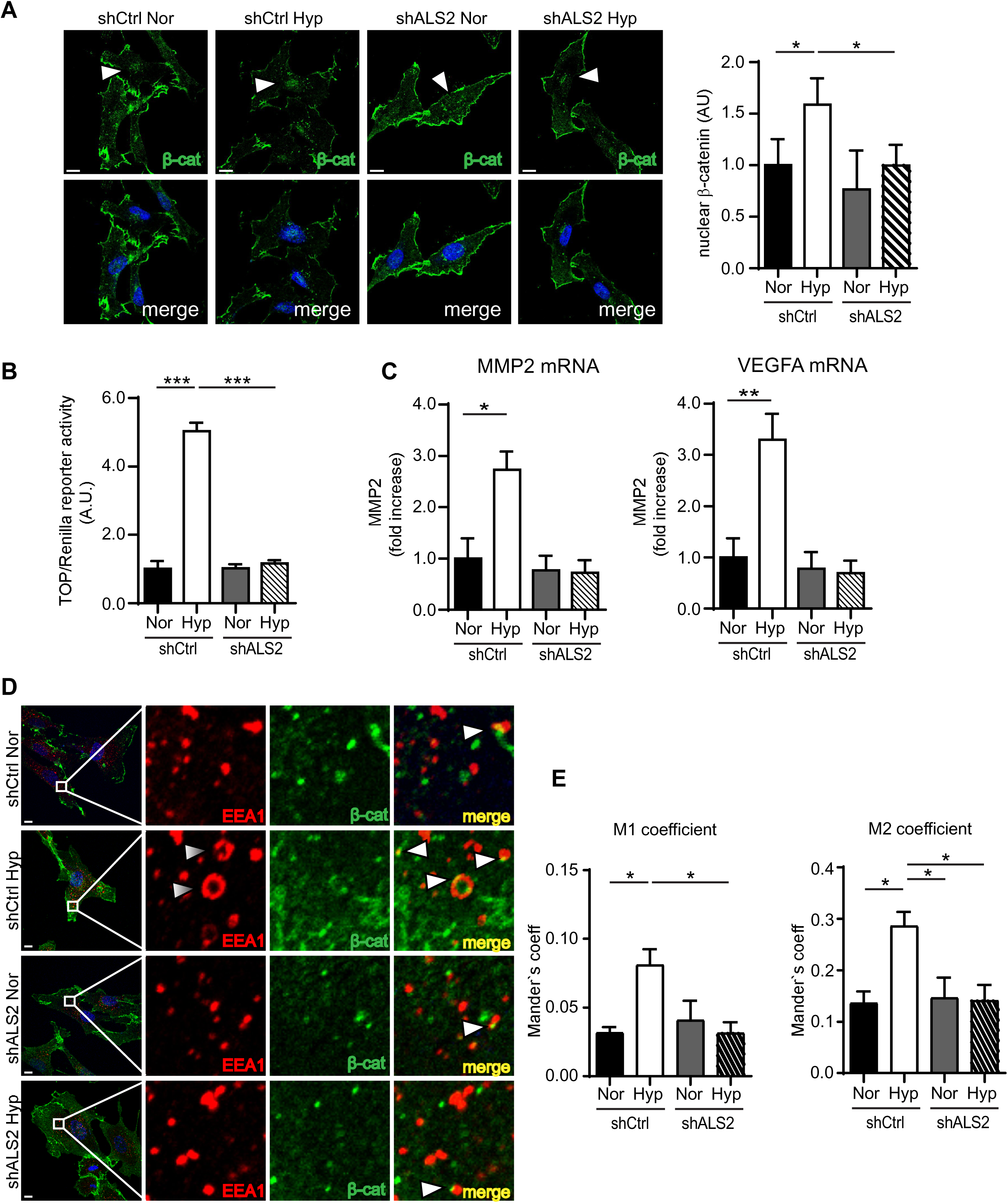
Requirement of ALS2 for sEVs-induced β-catenin activation in endothelial cells. **(A)** sEV fractions were prepared by differential centrifugation (100k pellets) from conditioned media obtained from A549 cells, treated with scramble (control) or ALS2 targeting shRNA, which were incubated for 24h in normoxia (Nor) or hypoxia (Hyp). Then, sEVs were added to normoxic EA.hy926 endothelial cells (1x10^4^ sEVs/cell), incubated for 24h and samples were fixed and stained with an antibody against total β-catenin, whereas nuclei were visualized with Hoescht. Representative confocal microscope images are shown. Bar represents 10 µm. Graph indicates the normalized nuclear fluorescence signal, obtained by dividing nuclear versus total fluorescence signal, as described in the materials and methods section, using the Image J software. Data represent the average from 3 independent experiments (mean±s.e.m.; *p<0.05). **(B)** EA.hy926 were co-transfected with the plasmid pTOP-FLASH (functional Tcf/Lef binding sites, black bars) along with the constitutively active vector encoding for renilla luciferase (as control), and then treated with sEVs obtained as in (A). After 24 h, cells were homogenized and prepared for Tcf/Lef reporter assays (see materials and methods for details). Data were obtained as the ratio luciferase/renilla activity and shown as the relative values with respect to non-treated cells. Data were averaged from 4 independent experiments (mean±s.e.m.; ***p<0.001). **(C)** EA.hy926 were stimulated for 24h with sEVs obtained as in (A) and then total RNA was extracted, for subsequent RT-qPCR analysis of MMP2 and VEGFA. Relative abundance of all screened mRNAs was normalized to actin, by using the ΔΔCt method. Graphs show data obtained by averaging 4 independent experiments (mean±s.e.m.; *p<0.05; **p<0.01). **(D)** EA.hy926 were treated with sEVs obtained as in (A), incubated for 24 h and then processed for immunofluorescence analysis of EEA1 and β-catenin. Representative confocal microscopy images and their magnifications are shown from 3 independent experiments (Bar 10µm). **(E)** Mander’s coefficients were determined from data in (D), using JACOP plugins of Image-J software, as described in the material and methods section. Data were averaged from 3 independent experiments (mean±s.e.m; *p<0.05). **(F)** Images described in (D) were analyzed for total and enlarged endosomes. Particle number per cell were analyzed (particles ˃0.05µm were scored). At least 5 images per condition were analyzed. The graph shows the average from 3 independent experiments (mean±s.e.m.;*p<0.05;**p<0.01). Enlarged/encircled structures were scored as positive events (at least 5 images per condition). The graph represents the average of 3 independent experiments (mean±s.e.m.; **p<0.01).

## DISCUSSION

### Vesicular transfer of ALS2

It has been largely known that the tumor microenvironment and specifically tumor hypoxia, promote the acquisition of aggressive traits, by inducing adaptative events, such as angiogenesis (36). The impact of hypoxia on tumor angiogenesis is associated with the modulation of endothelial cell responses via both direct and indirect events (5, 37, 38). Among indirect events associated with tumor hypoxia, the action of tumor cell-derived sEVs on endothelial cells remains to be better understood, especially regarding the identification of cargo that accounts for such effects (39). In this study, we showed that the guanine exchange factor (GEF) ALS2 is a novel sEV cargo, which was upregulated in hypoxia and that triggered a conserved signaling cascade in endothelial cells, accounting for pro-angiogenic events. In doing so, ALS2 was loaded within sEVs and transferred to recipient endothelial cells, promoting Rab5-GTP loading and hence downstream signaling via β-catenin, leading to endothelial cell migration and tube formation (see proposed model, Figure 8). To our knowledge, this is the first study reporting the consequences of transferring a vesicular GEF that accounts for GTPase activation in recipient cells. The concept of transferring any small GTPase regulators (i.e., GEFs or GAPs) within sEVs has not been explored before, although one recent study indicated that transferring of RNA regulators within sEVs leads to the upregulation of H1, a Rho-GEF that promotes RhoA activity in recipient colorectal cancer cells (40). However, as indicated, direct transferring of either GEFs or GAPs has not been documented for any small GTPase, including Rab proteins and hence, this is the first study reporting such mechanism.

**Figure 8.**
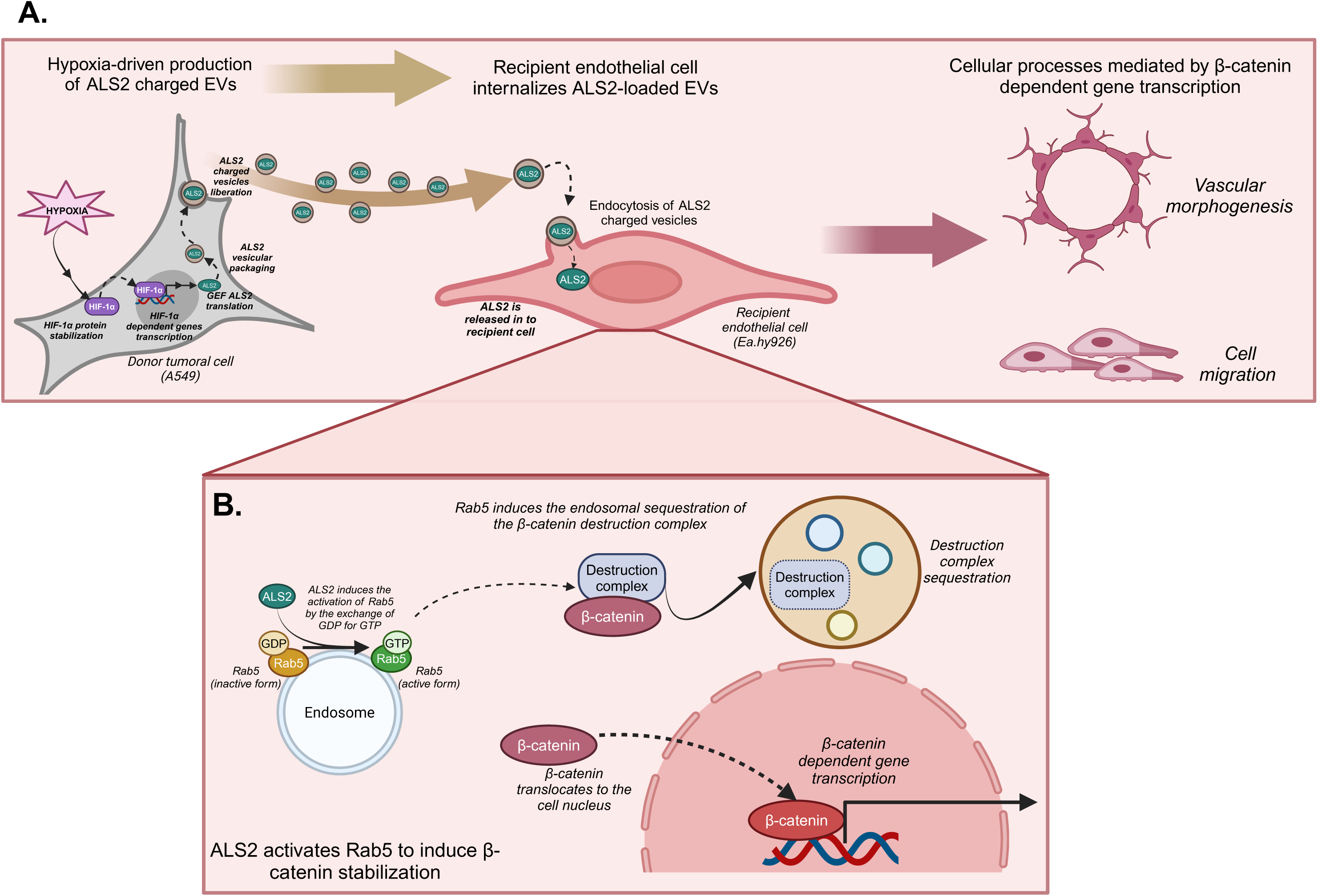
Proposed model. The scheme describes a model whereby sEVs released by hypoxic tumor cells promote angiogenic-related responses. Under hypoxic conditions, sEVs are liberated with the capacity of increasing endothelial cell migration and tube formation, via vesicular transfer of ALS2, which in turns activates Rab5-dependent signaling in endothelial cells, by promoting β-catenin signaling.

Previous studies showed that hypoxia-dependent effects in tumor cells are mediated by a signaling cascade involving stabilization of the hypoxia inducible factor HIF-1α, which via promoting transcription of ALS2, leads to the activation of the small GTPase Rab5 (23, 24). This signaling axis was relevant to the induction of tumor cell migration and invasion under hypoxic conditions, as shown in a wide panel of tumor cell models (24). Although all three proteins were upregulated by hypoxia in donor A549 tumor cells, hypoxia-induced loading within sEVs was evident for ALS2, but not for Rab5 and, most importantly, no changes in total Rab5 or HIF-1α were observed in recipient endothelial cells, thereby excluding the possibility that Rab5-dependent events were due to fluctuations in total levels of these proteins. In this context, although Rab5 and other Rab proteins are known to be incorporated within sEVs (41), no reports have addressed the effects of hypoxia on Rab composition within sEVs. Our study indicates that hypoxia does not affect Rab5 loading in sEVs, neither at total nor GTP-bound levels. Likewise, despite HIF-1α was previously shown to be incorporated as sEV cargo in donor tumor cells (42), we failed to notice any increases in recipient cell levels of HIF-1α (data not shown), further excluding the possibility that Rab5-GTP fluctuations were due to direct transfer of HIF-1α.

At the sEV-donor cell level, despite it is known that interfering with the endolysosomal pathway alters the bioactivity of cargo contained within sEVs (43), while other studies suggest that Rab5 is associated with exosome secretion in hepatocellular carcinoma and breast carcinoma cells (44, 45), we did not detect any significant fluctuations in particle size in ALS2-knockdown cells, which suggest that Rab5 activity was dispensable for these characteristics in this tumor cell model (lung carcinoma).

### Signaling in sEV-recipient endothelial cells

Endothelial cells are known to respond to different extracellular cues via conserved signaling pathways that converge in the induction of endothelial cell adhesion, migration and angiogenesis (46). Previous evidence indicates that signaling from Rab5-containing endosomal compartments is relevant for endothelial cell migration and angiogenesis *in vitro* (25, 26).

While sEVs are known to activate the Wnt/β-catenin pathway in recipient endothelial cells (18, 28), the possibility that these events are further modulated in hypoxic conditions remained unexplored. Here, we showed that hypoxia promotes the release of sEVs with increased capacity to stabilize and activate β-catenin-dependent events in endothelial cells and, most importantly, that these events depended on Rab5. While previous evidence obtained in epithelial cells indicates a direct relationship between Rab5 and β-catenin, via endosomal sequestration of the destruction complex (32), the feasibility of such regulation mechanism in endothelial cells was not explored before. Here, we established for the first time that the endosomal sequestration of the β-catenin destruction complex accounts for β-catenin proangiogenic events in endothelial cells. Besides our data, sEVs transfer of regulators of the Wnt/β-catenin pathway was previously documented (18) and hence, we cannot exclude the possibility that such transfer to recipient endothelial cells is operative in hypoxic conditions.

In summary, this study identifies a novel mechanism whereby tumor-derived sEVs, released in hypoxic conditions, promote proangiogenic-related responses in endothelial cells, via direct transfer of the Rab5GEF ALS2, which in turn activates Rab5 in recipient endothelial cells, leading to downstream signaling via β-catenin and hence increasing endothelial cell migration and angiogenesis *in vitro*. These findings will provide insights to better understand the role of the tumor microenvironment on angiogenesis associated with tumor progression.

## MATERIALS AND METHODS

### Antibodies

The following antibodies were used: anti-ALIX (sc-53540), anti-Axin (sc-293190), anti-Rab5 (sc-46692), anti-EEA1 (sc-33585), anti-α-Tubulin (sc-8035), anti-Actin (sc-47778), were from Santa Cruz Biotechnology (Dallas, TX, USA); anti-β-catenin (M3539) was from DAKO (DAKO, USA); anti-ALS2 (ab170896), anti-lamin A/C (ab108595), were from Abcam (Abcam, USA); anti-non-phosphorylated β-catenin (Ser33/37/Thr41 88145), anti-CD63 (52090), were from Cell Signaling Technology, Danvers, MA, USA; anti-GSK3β (610201), anti-GM130 (610823) from BD Bioscience, Becton, NJ, USA; anti-syntenin 1 (NBP2-76873), were from Novus Biologicals, Centennial CO, USA). Alexa Fluor 488 and Alexa Fluor 568 secondary antibodies were from Thermo Fisher Scientific (Waltham, MA, USA). Tissue culture medium, antibiotics, and fetal bovine serum were from Thermo Fisher and HyClone Laboratories (Logan, UT, USA). Glutathione– sepharose 4B was from GE Healthcare (Piscataway, NJ, USA. Control small interfering RNA (siRNA) sequence (sc-37007) and β-catenin-specific siRNAs (a pool of 3 siRNA sequences targeting human β-catenin (sc-29209), were from Santa Cruz Biotechnology. The EZ-ECL chemiluminescent substrate was from Pierce (Rockford, IL, USA). Wnt-C59 is the inhibitor of porcupine (inhibitor II C59) and was from Calbiochem (USA).

### Cell Culture

A549 lung carcinoma and EA.hy926 endothelial cells were cultured in DMEM-high glucose supplemented with 10% fetal bovine serum and antibiotics, according to the manufacturer’s instructions (American Type Culture Collection, Manassas, VA, USA). For transient transfection with plasmids, cells were grown for 24 h in complete medium at 50 to 70% confluence. Transfections were performed with 1 µg of the indicated plasmids (6-well plates) by using the Lipofectamine 2000 system (Thermo Fisher Scientific). For transient transfection with siRNA, cells were grown for 24 h in complete medium at 50 to 70% confluence. Transfections were performed with 10 nM of siRNA sequences (6-well plates) by using the Lipofectamine RNAiMax system (Thermo Fisher Scientific). Hypoxia was performed as previously described (24), using a hermetically sealed, modular incubator chamber (MIC-101 Billups-Rothenberg Inc., Del Mar, CA), and a gas mixture (1% O2, 5% CO2 and 94% N2, Linde Group).

### Small extracellular vesicles isolation

For small extracellular vesicles (sEVs) isolation, cells were cultured for 72 h in RPMI 1640 medium supplemented with 1% pen-strep and 5% exosome-depleted FBS (GibcoTM, ThermoFisher Scientific). sEV derived from A549 cells were isolated from the conditioned medium by differential ultracentrifugation, as previously described (27), using the P70AT fixed angle rotor (Hitachi-Himac CP100WX, Japan). Conditioned medium was centrifuged at: 300× g for 10 min 4 °C, (ii) 2000× g for 20 min 4 °C, (iii) 10,000× g for 35 min 4 °C, and (iv) 100,000× g for 75 min at 4 °C. After each step, the pellet was discarded, and the supernatant was used for the following step. The resulting pellet after the fourth step was resuspended in PBS and washed at 100,000× g for 75 min 4 °C to remove any contaminating proteins. The pellet was then resuspended in PBS.

### Nanotracking particle analysis

The size distribution and concentration of sEVs were analyzed by nanotracking particle analysis, as previously described (27). Nanotracking particle analysis was performed using NanoSight NS300 equipment (Malvern Instruments, Malvern, United Kingdom). The measurement was performed using the 532-nm laser and 565-nm long pass filter, with a camera level at 12 and a detection threshold of 4. sEVs were diluted in 1X PBS before the analysis.

### Western blot analysis

Cells were washed twice with cold PBS and lysed in 0.2 mM HEPES (pH 7.4) buffer containing 0.1% SDS, phosphatase inhibitors (1 mM Na_3_VO_4_), and a protease inhibitor cocktail. Total protein extracts (50 mg per lane, unless indicated) were separated by SDS-PAGE and transferred to nitrocellulose membrane. Blots were blocked with 5% gelatin in 0.1% Tween-PBS and then probed with antibodies. Bound primary antibodies were detected with horseradish peroxidase–conjugated secondary antibodies and the EZ-ECL system. sEVs markers were identified by Western blot. sEVs lysis was performed with cOmplete™ Lysis-M buffer (Roche Diagnostics GmbH, Mannheim, Germany), 10 µg of sEVs separated by SDS-PAGE gels were transferred to polyvinylidene difluoride membrane (PVDF; ThermoFisher Scientific) for 1 h at 100 V. Further, the membrane was analyzed by enhanced chemiluminescence with image capture on an iBright CL 1500 Imaging System (Thermo Fisher Scientific) (10.3390/ijms20204972).

### Plasmids

The pEGFP-C1 plasmids encoding Rab5/S34N (GDP-bound inactive mutant forms, negative dominant of Rab5) and ALS2ΔVPS (a dominant negative form of ALS2, which lacks GEF activity towards Rab5ALS2 have been previously described (23).

### Immunofluorescence

Immunofluorescence was performed as previously described (32). β-catenin, Axin, EEA1 and Rab5 were detected using their respective secondary antibodies conjugated to either Alexa Fluor 488 or Alexa Fluor 568. Samples were analyzed with a confocal microscope (LSM-Pascal 5; Carl Zeiss GmbH, Jena, Germany). β-catenin nuclear signal, analysis of particles EEA1 positives or Rab5 positives, and colocalization data were evaluated in original images, obtained by Confocal Microscopy and analyses were performed with the ImageJ software and the JACoP plugin, specifically for colocalization (Image Processing and Analysis in Java; National Institutes of Health, Bethesda, MD, USA; http://imagej.nih.gov/).

### Subcellular fractionation

Cells were washed twice with ice-cold PBS and homogenized in fractionation buffer supplemented with protease inhibitors and phosphatases. The cell homogenate was shaken at 30-50 rpm, for 30 minutes at 4°C. Subsequently, samples were centrifuged at 720g, 4°C for 5 minutes. The pellet was separated from the supernatant, to obtain nuclear and cytoplasmic proteins. The pellet was then washed with fractionation buffer and centrifuged at 720 g, 4°C for 10 minutes, removing the supernatant, and resuspending the pellet in lysis buffer.

### β-catenin Tcf/Lef reporter assay

For Tcf/Lef transcriptional activity measurements, the luciferase reporter activity system was used, as previously described (32). This assay is based on co-transfection with the plasmid pTOP-FLASH, which encodes for the luciferase gene preceded by three tandem Tcf/Lef binding sites, and the constitutively active vector encoding for Renilla luciferase. Cells were transfected with 1 μg (TOP) and 0.25 μg (Renilla), for 24 h, and then cells were homogenized according to the manufactureŕs instructions (Dual-Luciferase resporter assay system, Promega, Madison, WI, USA)

### RNA extraction and analysis

Total RNA was extracted with TRIZOL (Invitrogen, Life Technologies). For complementary DNA (cDNA) synthesis, samples were treated with RNase-Free DNase Kit (#M6101; Promega), quantified, and purity was verified for subsequent reverse transcription, using the cDNA Reverse Transcription Kit (Applied Biosystems). MMP2, VEGFA and actin were quantified by quantitative reverse transcription polymerase chain reaction (ΔΔCt method; Applied Biosystems), using the following forward and reverse primers, respectively: VEGFA, 5′-CTC TAC CTC CAC CAT GCC AAG-3′, 5′-AGA CAT CCA TGA ACT TCA CCA CTT-3′; MMP2, 5′-GAT ACC CCT TTG ACG GTA AGG A-3′, 5′-CCT TCT CCC AAG GTC CAT AGC-3′ and actin, 5′-TGG CAC CCA GCA CAA TGA AGA-3′, 5′-GAA GCA TTT GCG GTG GAC GAT-3′.

### Proteinase-K protection assay

Proteinase-K protection assays were performed as previously described (33). Briefly, cells were harvested from 10 cm plates, collected by centrifugation at 4°C, and then resuspended in buffer (100 mM potassium phosphate pH 6.7, 5 mM MgCl_2_, 250 mM sucrose) containing 6.5 mg/mL digitonin, incubating them for 5 minutes at room temperature, followed by 30 minutes on ice. Digitonin solution was removed by centrifugation at 13000 rpm (5 minutes) in an eppendorf centrifuge. Permeabilized cells were resuspended in buffer without digitonin and aliquoted into eppendorf tubes containing stocks of reagents that will generate final concentrations of 1 mg/mL Proteinase K, 1 mg/mL Proteinase K with 0.1% Triton X-100, or nanopure water. Samples were incubated for 10 minutes at room temperature and the reaction was stopped by the addition of pre-heated loading buffer containing 20 mM PMSF, followed by an additional heating at 95°C (5 minutes). Samples were analyzed by SDS-PAGE and Western blotting.

### Rab5-GTP pull-down assay

Rab5-GTP levels were measured using the active Rab5 binding domain (GST-R5BD) in pulldown assays, as previously described (24).

### Migration assays

Cell migration was evaluated in wound healing and Boyden chamber assays (Transwell Costar, 6.5 mm diameter, 8 µm pore size), as previously reported (26).

### *In vitro* vascular network formation assay

The *in vitro* tubule formation assay was performed as previously described (26). To this end, EA.hy926 cells were seeded on top of growth factor–reduced Matrigel-coated plates (3x10^4^ cells per well in serum-free medium) and incubated for 6 to 8 h at 37°C. Cell cultures were photographed (5 pictures per condition) and data were analyzed by ImageJ software (Image Processing and Analysis in Java; National Institutes of Health, Bethesda, MD, USA; http://imagej.nih.gov/).

### Construction plasmid encoding shRNAs

Short hairpin RNAs (shRNA) were cloned (Genscript) into HpaI and XhoI downstream of the U6 promoter of the self-inactivating lentiviral vector pLL3.7 that was modified replacing the EGFP sequence by puromycin sequence (pLL3.7-Puro). The target sequence of ALS2 used was sh-ALS2 5’-CTGTCAGATCATTCAGTAAGAGA-3. A control shRNA directed to Luciferase (sh-Ctrl) was generated using the target sequence 5’-TTCTCCGAACGTGTCACGT-3′.

### Bacterial transformation and plasmid DNA isolation

DH5α (InvitrogenTM, ThermoFisher Scientific, Carlsbad, CA, USA) for pCMV-VSV-g, and pS-PAX2 vectors, and One ShotTM Stbl3TM *E. coli* (InvitrogenTM, ThermoFisher Scientific, Carlsbad, CA, USA) for pLL3.7-Puro, were transformed with the plasmids, following manufacturer’s instructions. The transformed bacteria were grown to plasmid DNA isolation with NucleoBond Xtra Midi kit (Machery-Nagel, Allentown, PA, USA) according to the manufacturer’s protocol. The plasmid DNA isolated was quantified using the spectrophotometer EPOCH2 microplate reader (BioTek, Winooski, VT, USA).

### Generation of stable knockdown cell lines with shRNA lentiviral particles

Lentiviral particles were generated by co-transfection of HEK293T cells with pLL3.7-shRNA, pCMV-VSV-g, and pS-PAX2 vectors, harvested and concentrated as previously described (47). After 60 h, the resulting supernatant was harvested, centrifuged at 100 x g for 5 min, and filtered through 0,45 µm cellulose acetate filters to eliminate cell debris. The lentiviral particles were centrifugated for 1 hour 30 min at 100,000 x g in a P28S swinging bucket rotor (Hitachi-Himac CP100WX, Japan). The supernatant was discarded, and the particles were resuspended in 200 µL.

### Cell transduction

Silencing of ALS2 was generated by transduction with shRNA-containing lentiviral particles and a shRNA against luciferase was used as a control. For transduction, A549 cells were seeded at 2x10^4^ cells/well into 12-well plate. At 8 h after seeding, cells were transduced with shRNA-containing lentiviral particles for 72 h, and 48 h post-transduction were selected and expanded with 4 μg/mL puromycin (GibcoTM, ThermoFisher Scientific).

### Statistical analysis

Where pertinent, results were compared by Student’s t-test with Welch’s correction, or 2-way ANOVA by GraphPad Prism 5 software (GraphPad Software, La Jolla, CA, USA). Values averaged from at least 3 independent experiments were compared (average ± SEM, standard error of the mean). A value of p<0.05 was considered significant.

## Supporting information

Cover Letter

## ACKNOWLEDGMENTS

This work was supported by the National Fund for Scientific and Technological Development, FONDECYT 1220517 (to V.T.); FONDECYT 1230983 (to M.V.); the Advanced Center for Chronic Diseases, FONDAP-ACCDiS 15130011 (to V.T. and M.V.); Millennium Institute on Immunology and Immunotherapy ICN09_016/ICN 2021_045 (to V.T.); ANID Fellowship 21230341 (to B.G). We acknowledge Ana Riveros Salvatierra (Faculty of Chemical and Pharmaceutical Sciences, Universidad de Chile) for her technical assistance on particle analysis determination.

## AUTHOR CONTRIBUTIONS

Torres and M. Varas designed research, performed research, contributed with reagents and analytic tools, analyzed data, and wrote the paper; P. Silva designed research, performed research, analyzed data, and reviewed the paper; N. Hernández, H. Tapia, B. Gaete, T. Flores, D. Herrera and A. Cáceres performed research, analyzed data and reviewed the paper.

