## Supplementary Figures for "Tumor-derived hypoxic small extracellular vesicles promote endothelial cell migration and tube formation via ALS2/Rab5/β-catenin signaling"

### SUPPLEMENTARY FIGURE 1

**A**

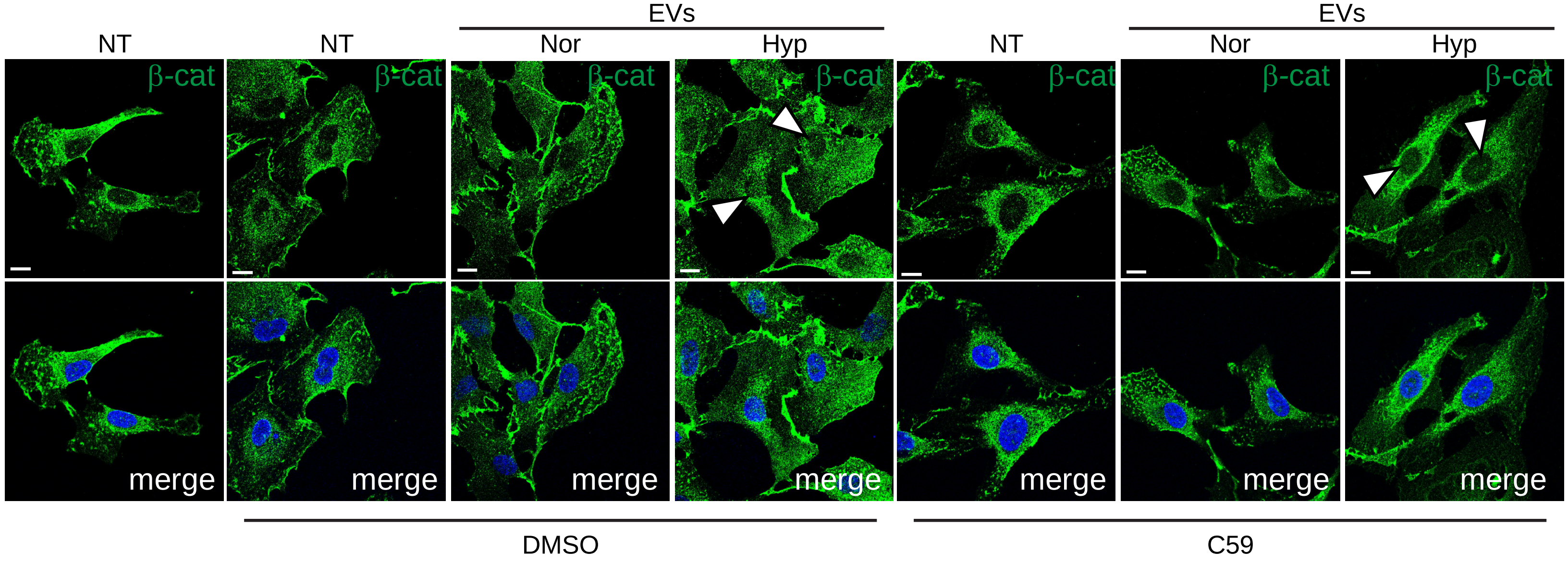


**B**

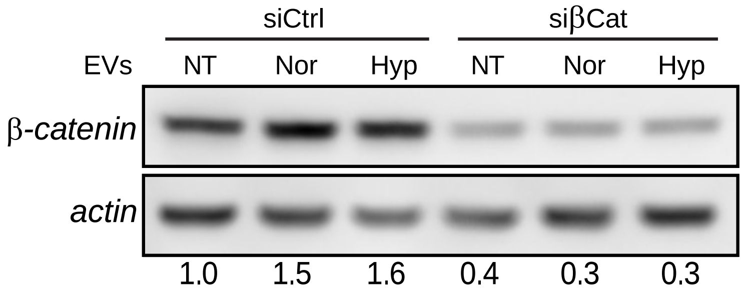


**(A)** EA.hy926 cells were incubated or not with normoxic (Nor) or hypoxic (Hyp) EVs for 24h, as described in the materials and methods, in the presence of vehicle (DMSO) or C59 (5 µM). Then, samples were fixed and stained with an antibody against total β-catenin, whereas nuclei were visualized with Hoescht. Representative confocal microscope images are shown. Bar represents 10 µm. **(B)** EA.hy926 cells were transfected with a pool of siRNA constructs targeting luciferase (siCtrl) or β-catenin (siβcat) and then incubated or not with normoxic (Nor) or hypoxic (Hyp) EVs for 24h. Total protein extracts were obtained and analyzed for subsequent Western blot of β-catenin and actin. Representative images are shown and numbers below panels indicate the scanning densitometric analysis.

### SUPPLEMENTARY FIGURE 2

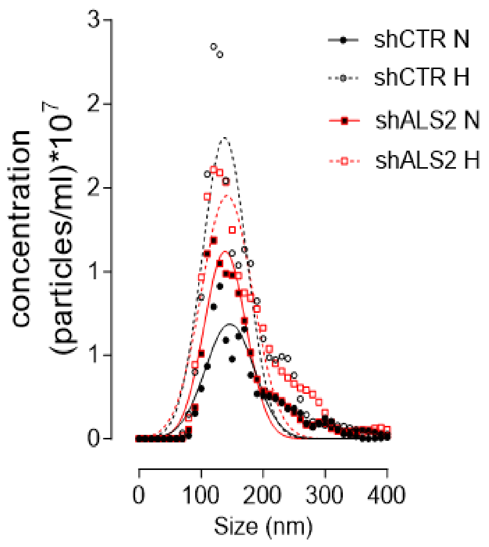


Particle size analysis. A549 cells stably expressing either control or ALS2-targeting shRNA constructs were cultured in exosome-free medium, for 24h in normoxia (Nor) or hypoxia (Hyp). Conditioned media were obtained and EV fractions were prepared by differential centrifugation. EV fractions (100k pellets) were analyzed for size distribution and concentration in a NanoSight NS300 (Malvern).
