## Supplementary material for "Tumor-derived hypoxic small extracellular vesicles promote endothelial cell migration and tube formation via ALS2/Rab5/β-catenin signaling": Cover Letter

FACULTAD DE
ODONTOLOGÍA
UNIVERSIDAD DE CHILE

*January 31st, 2024*
*Santiago, Chile*

*The FASEB Journal*

*Dear Editor,*

We are submitting our study entitled "Tumor-derived hypoxic extracellular vesicles promote endothelial cell migration and tube formation via ALS2/Rab5/β-catenin signaling", by the authors Patricio Silva, Nadia Hernández, Héctor Tapia, Belén Gaete, Tania Flores, Daniela Herrera, Albano Cáceres, Manuel Varas-Godoy and Vicente A. Torres, for consideration in *The FASEB Journal*.

Tumor hypoxia is associated with cancer progression, angiogenesis and metastasis via modifications in the release and cargo composition of extracellular vesicles (EVs) secreted by tumor cells. Specifically, hypoxia-induced release of EVs is known to promote tumor angiogenesis by modulating the function of endothelial cells via mechanisms that remain to be better understood, especially regarding the identification of cargo that accounts for such effects. In this study, we identify the guanine exchange factor ALS2 as a novel EV cargo, which is upregulated in hypoxia and that triggers a conserved signaling cascade in endothelial cells, accounting for pro-angiogenic events. In doing so, ALS2 is loaded within EVs and transferred to recipient endothelial cells, activating the small GTPase Rab5 and hence downstream signaling via β-catenin, leading to endothelial cell migration and angiogenesis *in vitro*. This is the first study reporting the consequences of transferring a small GTPase regulator to recipient endothelial cells via EVs. Therefore, these findings open new avenues to understand the mechanisms whereby hypoxia and the tumor microenvironment modulate tumor angiogenesis.

In conclusion, our results reveal a novel mechanism whereby tumor-derived EVs, released under hypoxic conditions, promote endothelial cell responses associated with tumor angiogenesis. We believe that our results will be of interest to a broad range of researchers working in the areas of cell biology, cancer research, and signal transduction, and especially to the readership of *The FASEB Journal*. The authors declare that there are no conflicts of interest.

Please address all correspondence to:
Vicente A. Torres, Associate Professor
Institute for Research in Dental Sciences
Faculty of Dentistry
Universidad de Chile
Olivos 943, Independencia, Santiago, Chile


Dr. Manuel Varas-Godoy , Associate Professor
Centro de Biología Celular y Biomedicina
Facultad de Medicina y Ciencia
Universidad San Sebastián
Carmen Sylva 2444 Santiago, Chile
